# No Evidence in Favour of the Existence of ‘Intentional’ Binding

**DOI:** 10.1101/2023.02.06.526214

**Authors:** Gaiqing Kong, Cheryne Aberkane, Clément Desoche, Alessandro Farnè, Marine Vernet

## Abstract

Intentional binding refers to the subjective temporal compression between a voluntary action and its sensory outcome. Despite some studies having challenged the link between temporal compression and intentional action, the intentional binding is still widely used as an implicit measure for the sense of agency. The debate remains unsettled primarily because the experimental conditions used in previous studies were confounded with various alternative causes for temporal compression, and action intention has not yet been tested comprehensively against all potential alternative causes in a single study. Here, we solve this puzzle by jointly comparing participants’ estimates of the interval between three types of triggering events with comparable predictability - voluntary movement, passive movement, external sensory event - and an external sensory outcome (auditory or visual across experiments). Results failed to show intentional binding, i.e., no shorter interval estimation for the voluntary than the passive movement conditions. Instead, we observed temporal (but not intentional) binding when comparing both movement conditions to the external sensory condition. Thus, temporal binding seems to originate from sensory integration and temporal prediction, not from action intention. As such, these findings underscore the need to reconsider the use of "intentional binding" as a reliable proxy of the sense of agency.

**Public Significance Statement:** When we press a light switch and observe a bulb lightening, we experience a sense of agency, a feeling of control over these events. We also estimate the temporal interval between our voluntary action and its consequence shorter than the same interval between two events in which we are not involved. Such temporal binding has thus been taken as a measure of the sense of agency. However, our study reveals that voluntary actions are neither necessary, nor sufficient for temporal binding. Instead, temporal binding relies on predicting and integrating information. The sense of agency can be disturbed in various psychiatric disorders and its brain mechanisms are currently actively explored. Our study urges amending how it is measured.

## Introduction

The intentional binding effect (also sometimes called temporal binding) refers to the subjective compression of the perceived temporal interval between a voluntary action and its sensory consequence (Haggard et al., 2002; Moore & Obhi, 2012). It has been widely used as an implicit measure for the Sense of Agency (SoA), the human feeling of controlling one’s own actions and, through them, their consequences in the external world (Haggard & Chambon, 2012).

There are two well-established paradigms to measure intentional binding: the single event time estimation procedure, with a variant of the Libet clock paradigm (Borhani et al., 2017; Desantis et al., 2012; Engbert & Wohlschläger, 2007; Haggard et al., 2002; Kirsch et al., 2019; Kong et al., 2017; Ruess, Thomaschke, Haering, et al., 2018; Tsakiris & Haggard, 2003), and the interval estimation procedure (Buehner & Humphreys, 2009; Caspar et al., 2021; Caspar et al., 2020; Engbert et al., 2008; Engbert et al., 2007; Ohata et al., 2022; Poonian & Cunnington, 2013; Suzuki et al., 2019; Zapparoli et al., 2020). In the Libet clock variant, participants make a self-paced button-press action that triggers a tone (usually 250 ms later) while viewing a small rotating clock hand. It is commonly reported that participants experience a sense of agency over the tone. Haggard et al. (2002) showed that voluntary actions, in contrast to involuntary ones, elicit a binding effect such that the perceived time of the action is delayed (biased towards the outcome) while the perceived time of the outcome is advanced (biased towards the action). In the interval estimation task, participants are typically required to give an explicit numerical estimate of the interval between an action and its outcome (commonly a tone), which are separated by a short but variable delay. Similarly to the former paradigm, participants judge the interval as being shorter in the voluntary condition relative to a baseline, which can be the interval between two successive external events (Buehner & Humphreys, 2009; Dewey & Knoblich, 2014; Imaizumi & Tanno, 2019), or the interval between an involuntary action and its outcome (Caspar et al., 2015; Engbert et al., 2008; Zapparoli et al., 2020).

Over the years, studies on intentional binding have yielded conflicting evidence. On one hand, temporal compression similar to the intentional binding was also observed when action intention was absent (Buehner, 2012; Gutzeit et al., 2023; Kong et al., 2017; Suzuki et al., 2019), or when participants did not move, but simply observed a movement on a video (Poonian & Cunnington, 2013). Additionally, Buehner (2015) replicated the original Haggard et al. (2002) study with the Libet clock paradigm, but replaced the TMS-induced involuntary muscle twitch with a machine-induced involuntary key press. The results showed a reduced, but still significant binding effect for an involuntary causal movement condition. These studies thus suggest that one’s own intention is not necessary for eliciting temporal binding. On the other hand, Buehner & Humphreys (2009) showed that intentional action that is not causal is not sufficient for temporal binding to occur, and a recent study found no difference in temporal estimation between voluntary and involuntary actions when the temporal predictability of events was equalized between conditions (Kirsch et al., 2019), further suggesting that action intention alone is not sufficient either to elicit temporal binding effects.

A systematic review revealed that the effect size of temporal binding depends heavily on the condition used as baseline (Tanaka et al., 2019). This could be ascribed to some confounding variables, as action intentionality was not always the only feature varying between the operant condition (involving a voluntary movement) and the baseline. For example, in cases when an external sensory event was used as the first event at baseline, participants merely observed (or listened to) two external events. In such cases, action was not involved in the baseline and participants had no access to movement-related information, including voluntary motor command and somatosensory feedback that is known to influence time perception (Cao et al., 2020; Hagura et al., 2012; Tomassini et al., 2014; Wiener et al., 2019). Thus, to isolate the “intentional” character of intentional binding, a passive movement condition can serve as a better baseline. This baseline, whereby an involuntary movement is induced either mechanically or by the experimenter, was absent in many previous intentional binding studies (Suzuki et al., 2019). Furthermore, the comparison of these two baselines (external sensory event and passive movement as the first event), which is crucial to disentangle the role of somatosensory information during action execution from that of intention in temporal binding, has only been explored by Buehner (2015) and Kirsch et al. (2019). Kirsch et al. (2019) compared, with the Libet paradigm, voluntary and involuntary movements to an external event as a baseline, but in this study the baseline was a single event (a tone). Thus, there was a different number of events in the operant condition (two events: voluntary movement and a tone) and the baseline (one event only: a tone). As interval segmentation by events is known to influence time perception (Bangert et al., 2020; Faber & Gennari, 2017), the different number of events could introduce an additional confounding variable. Buehner (2015) aimed to test the role of perceived causality between the first and second events and therefore equalized it in both the active and passive movement conditions. However, the predictability of the first event was not controlled for across conditions, another potential contributor to bindings. Indeed, in addition to the type of baseline, another possible confounding component is the predictability of the first event (Buehner, 2012; Gentsch et al., 2012; Hughes et al., 2013; Suzuki et al., 2019). When previous studies used involuntary/passive movements as baselines, their onset remained unpredictable for participants (Caspar et al., 2015; Engbert et al., 2008; Zapparoli et al., 2020). To our knowledge, only two studies controlled for the predictability of the movement while comparing active and passive movement conditions. One controlled for the temporal predictability of passive key-presses with the method of constant stimuli (i.e., comparing the duration of an interval to the duration of a tone), and reported that intentional binding was observed for 600-ms intervals, but not for 250-ms intervals (Nolden et al., 2012). However, passive movements came from the key-board popping the participants’ fingers upward, thus involving different somatosensory feedback compared to active movements. The other study that controlled for the temporal predictability of the passive movements actually reported no difference in temporal estimation between voluntary and involuntary movements (Kirsch et al., 2019). However, this study applied single event time estimation with the Libet clock paradigm, which could differ from the binding effect as measured with the interval estimation task. Indeed, previous studies found that the binding effect increases with increasing intervals when the interval estimation method is applied, but that it decreases when the Libet clock method is used (Buehner & Humphreys, 2009; Haggard et al., 2002). Thus, the second aim of the present study was to test whether binding effects emerge when the temporal predictability of the first event is controlled for, using the interval estimation paradigm.

Finally, the sensory modality of the action’s consequence (outcome) could also influence the binding effect. While Engbert and colleagues (2008) found comparable amounts of the binding effect across auditory, somatic and visual modalities, another study reported that the overall intentional binding effect is weaker with visual than with auditory outcomes (Ruess, Thomaschke, & Kiesel, 2018). Thus, the third aim of the present study was to compare binding effects between visual and auditory outcomes.

To summarize, we aimed to tease apart the roles played by action intention, temporal predictability and somatosensory information in temporal binding. To this aim, we measured the magnitude of temporal binding with the interval estimation procedure by comparing an active movement condition to two different baselines, namely, the passive movement and the external sensory event conditions. In addition, we controlled for the temporal predictability of the first event. Finally, in order to assess the generality of the action effect across sensory modalities, the outcome was either auditory (Experiment 1) or visual (Experiment 2). Intentional binding should be demonstrated by shorter interval estimation for the active than the passive movement condition. Crucially, other types (i.e., not intentional) of temporal binding or dilation could be shown by comparing different interval estimations, namely the passive movement condition and the external event condition. Indeed, in such a comparison, intention is entirely absent.

## Experiment 1

### Method

#### Transparency and Openness

We report how we determined our sample size, data exclusions, manipulations, and experimental measures, and follow Journal Article Reporting Standards (JARS) (Kazak, 2018). Data, task, and materials for all experiments are available at the project’s Open Science Framework page (https://osf.io/n7y8a/). Data were analyzed using Python, Version 3.0. The study’s design and analysis were not preregistered.

#### Participants

For Experiment 1 (auditory action outcome), we initially recruited 24 participants to match the sample size of a previous study using a similar interval estimation procedure (Caspar et al., 2015) and ensure enough power according to this same study (effect size=0.938, desired power=0.95, required sample size=17). However, with this sample size, we did not find any difference between active movement and passive movement conditions. To definitively rule out that the lack of binding effect was due to lack of power, we conducted another power analysis based on a former study with estimated effect size of 0.593 (Engbert et al., 2008). This analysis suggested a required sample size of 39 participants, and we increased our sample size to 44 participants. This sequential design prevents us from interpreting any significant effect using standard statistical analyses. In order to assess the existence of binding, we instead relied on Bayes analyses in the rest of this study. One participant was excluded due to failure to produce temporal intervals varying monotonically with actual intervals. The remaining 43 participants (28 females, age = 25.2 ± 4.6 years old) participated in all three conditions.

Participants were naive as to the purpose of the study and reported normal or corrected-to-normal vision, normal audition and no neurological history. All participants provided informed consent before participation and received a payment for their participation. Procedures were approved by the ethics committee (CEEI/IRB Comité d’Evaluation Ethique de l’Inserm, n°21-772, IRB00003888, IORG0003254, FWA00005831) and adhered to the ethical standards of the Declaration of Helsinki except for registration in a database. Data was collected in 2021.

#### Apparatus and setup

The action consisted of a key-press performed on a keypad, which was placed at the left side of the monitor (resolution of 1280 × 1024) used to display a fixation cross (Figure 1A). The view of the keypad was prevented by an opaque board. The action outcome for the interval estimation task was an auditory tone (sampling rate: 44.1K Hz, 30 ms duration) played over a loudspeaker. Participants’ estimates were collected via another external keypad operated by their right hand. The experiment was programmed using the unity platform (Unity 2018.4.22f1 Personal) and Microsoft Visual Studio 2019.

**Figure 1.**
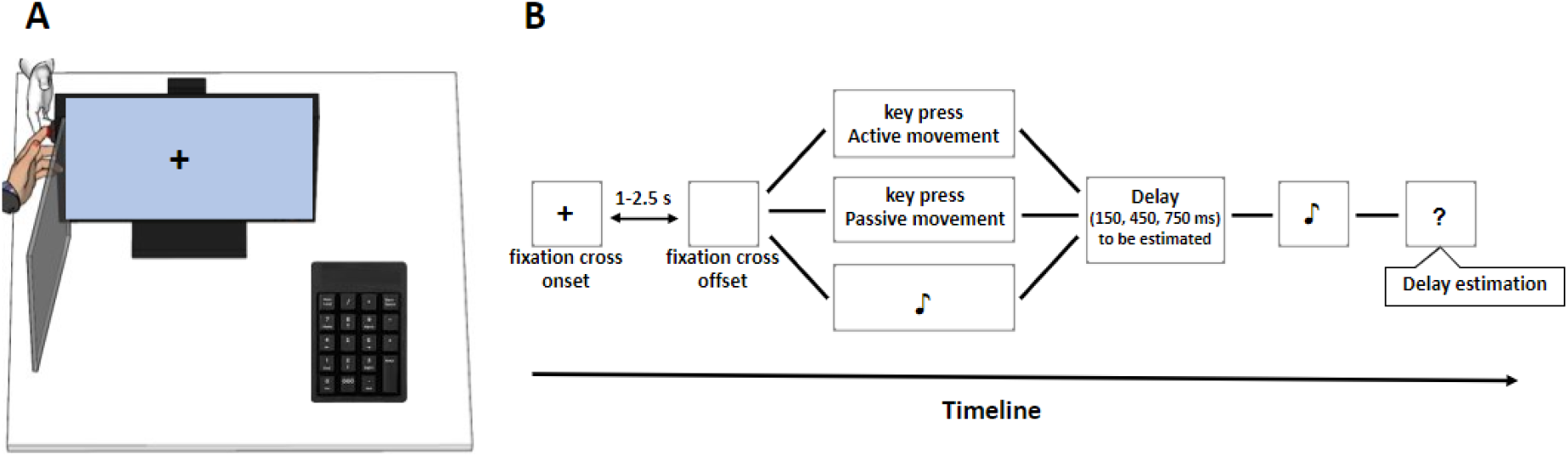
Set-up (A) and the three experimental conditions (B) of Experiment 1. **(A)** The set-up is shown here for the passive movement condition. **(B)** Schematic illustration of a trial in the three conditions: active movement, passive movement and external sensory event conditions. Note that the same number of events occurred during a trial for all the conditions.

#### Task and procedure

Participants sat on a chair and viewed the monitor at approximately 60 cm (Figure 1A). Prior to the experiment, participants were first invited to read the instructions, and the experimenter also provided verbal instructions during the practice phase. Three blocked conditions (active movement, passive moment, external sensory event) were then administered in counterbalanced order (Figure 1B). For all conditions, each trial started with a fixation cross, which was presented with a random duration between 1 s and 2.5 s. The fixation cross offset served as a temporal cue to signal the occurrence of the first event in all conditions: informing that key-press was allowed at one’s own willing time in the active movement condition; informing of the forthcoming passive key-press in the passive movement condition, where the experimenter pushed down the participant’s finger after the fixation cross offset, attempting to match the offset typically produced by the participants in a pilot experiment; or informing of the forthcoming first tone in the external sensory event condition, which was played 500 ms after the fixation cross offset. Due to a recording error, the duration between the offset of the fixation cross and the first event (“Fix-off/First-event”) was not directly recorded. Instead, the duration between the trial onset and the first event (“Trial-on/First-event”) was recorded. Given that the duration between the trial onset and the fixation cross offset (“Trial-on/Fix-off”) was uniformly randomized between 1 and 2.5 s across all conditions, a similar “Fix-off/First-event” duration across conditions could be inferred from a similar “Trial-on/First-event” duration across conditions. This was indeed the case for the active and passive movement conditions (on average 2.55 s for the active and 2.72 s for the passive movement conditions, Mann-Whitney U test, p = 0.07). Concerning the external event condition, the distribution of the “Trial-on/First-event” duration did not mirror the trial-to-trial variability observed with the active and passive movement conditions (on average 2.30 s), but allowed maintaining the predictability of the first event. A technical error in a subgroup of 12 participants led to different timings for the external event condition only: the “Trial-on/Fix-off” duration was uniformly randomized between 1 and 1.5 s (instead of 1 and 2.5 s) and the “Fix-off/First-event” was uniformly randomized between 0.5 and 1.5 s (instead of a fixed 0.5 s interval). In this subgroup, the predictability of the first event in the external event condition was lower. For this reason, all the analyses were run again without this subgroup, and all statistical results that are different from the main analyses are reported as footnotes in the manuscript. Hence, the disappearance of the fixation cross provided participants with similar levels of predictability for the first event in all conditions. Therefore, the three conditions differed mainly regarding the information available about the action. In the active movement condition, participants had predictability, intentional efferent and proprioceptive information about their own action. In the passive movement condition, they had predictability and proprioceptive, but lacked intentional efferent information about the action. In the external sensory event condition, no action related information was available to participants, but they had a similar level of predictability of the first event (Table 1).

**Table 1.** The three conditions and the available information about the first event.

| Information / First event | Action intention | Somatosensory information | Predictability |
| --- | --- | --- | --- |
| Active movement | ✓ | ✓ | ✓ |
| Passive movement | ✗ | ✓ | ✓ |
| External sensory event | ✗ | ✗ | ✓ |

In the active movement condition, participants performed a voluntary key-press with their left index finger whenever they wanted after the fixation cross disappeared, which generated a subsequent tone. In the passive movement condition, the experimenter, sitting behind the monitor and wearing a glove, pressed the participant’s passive left index finger down onto the button to generate the same tone (Caspar et al., 2015; Zapparoli et al., 2020). In the external sensory event condition, participants merely listened to one tone followed by a second occurrence of the same tone. The delay between the first event (active/passive keypress or first tone) and the second event (subsequent tone) was chosen pseudo-randomly among the following intervals: at 150, 450 or 750 ms, to make sure participants complied with the temporal estimation instruction (Figure 1B).

For active and passive movement conditions, participants were asked to estimate the elapsed interval (in ms) between the onset of the index finger action and the onset of the tone emission. For the external sensory event condition, they were asked to estimate the interval between the onset of the first and of the second tone. Participants were told that the delay varied randomly from trial to trial, and never exceeded 1000 ms. They were encouraged to use the full range between 1 and 1000 ms to express even slight variations in their experience of the time elapsed. During the practice phase, participants were reminded that 1 s would correspond to a judgment of 1000, 0.5 s would correspond to 500, and so forth. Participants practiced randomly with 10 different intervals from 100 ms to 1000 ms to have the impression that the delay varied on a trial-by-trial basis. During the formal testing, each interval (150, 450 and 750 ms) was randomly presented once for the first three trials, and then 21 times randomly, resulting in 66 trials per condition. The first three trials were discarded from analysis.

#### Data Analysis

An intentional binding should be demonstrated by the presence of shorter interval estimates for the active than the passive movement condition. Other types of temporal binding (i.e., not intentional) could be shown by the presence of different interval estimates between the active and/or passive movement conditions and the external event condition. To explore the existence or the absence of an effect of interest (namely intentional or non-intentional bindings), we calculated Bayes factors (BF) for the relevant paired comparisons. To calculate the BF for directional predictions of differences between the active movement condition and the two other conditions (passive movement and external sensory event), we used a half-normal distribution with a standard deviation of 122.5 ms, which was the size of the largest binding reported in a previous study (Caspar et al., 2015). For the comparison between the passive movement condition and external sensory event condition, we made no directional prediction and calculated a BF using a uniform distribution with a minimum of 13.5 ms, which was the smallest binding reported in the same study (Caspar et al., 2015), and a maximum of 122.5 ms (see also Suzuki et al., 2019). A BF of above 3 would indicate substantial evidence for the existence of a binding, a BF below 1/3 would indicates substantial evidence for the inexistence of a binding, and intermediate values would not provide any substantial evidence either way (Dienes, 2014; Dienes & Mclatchie, 2018; Jeffreys, 1939; Wagenmakers et al., 2017). We also reported the robustness region (RR), giving the range of scales (i.e., the minimum and maximum standard deviation of the half normal distribution, or the minimum and maximum difference between the higher bound and the lower bound of the uniform distribution), which would lead to the same conclusion (Dienes, 2019).

## Results

The evidence for an intentional binding, i.e., shorter mean interval estimation for the active movement condition (Mean ± SE = 286.93 ± 19.11) than the passive movement condition (Mean ± SE = 303.46 ± 19.71) was inconclusive (BF_HN(0,122.5)_ = 0.82, RR_1/3<B<3_ = [0, 307])^1^. The Robustness Regions indicate that even smaller predictions of the size of the intentional binding would lead to the same conclusion. On the contrary, the interval estimation for the active movement condition was shorter than that observed in the external sensory event condition (Mean ± SE = 335.74 ± 22.32), with substantial evidence in favor of the existence of such a temporal binding (BF_HN(0,122.5)_ = 60.6, RR_B>3_ = [6, 2693]). Finally, the evidence for shorter interval estimation for the passive movement condition than the external sensory event condition was inconclusive (BF_U(13.5,122.5)_ = 2.5, RR_1/3<B<3_ = [93, 829])^2^.

When considering actual delays separately, the evidence for an intentional binding, i.e., a difference between the active and passive movement conditions, was inconclusive across all intervals (long: BF_HN(0,122.5)_ = 0.85, RR_1/3<B<3_ = [0, 319]; middle: BF_HN(0,122.5)_ = 0.38, RR_1/3<B<3_ = [0, 141]; short: BF_HN(0,122.5)_ = 0.48, RR_1/3<B<3_ = [0, 175])^3^. On the contrary, the evidence for a non-intentional binding when comparing active movement vs. external sensory conditions was substantial for both the long and middle intervals (long: BF_HN(0,122.5)_ = 591, RR_B>3_ = [8, >3000]; middle: BF_HN(0,122.5)_ = 30, RR_B>3_ = [7, 1346]); for the short interval there was substantial evidence for an absence of binding (BF_HN(0,122.5)_ = 0.18, RR_B<1/3_ = [64, >3000]). Finally, the evidence for a non-intentional binding when comparing passive movement vs. external sensory conditions was substantial for the long interval (BF_U(13.5,122.5)_ = 80, RR_B>3_ = [0, >3000]), and the evidence was inconclusive for the middle and short intervals (middle: BF_U(13.5,122.5)_ = 2.86, RR_1/3<B<3_ = [104, 932]); short BF_U(13.5,122.5)_ = 0.88, RR_1/3<B<3_ = [30, 286])^4^.

**Figure 2.**
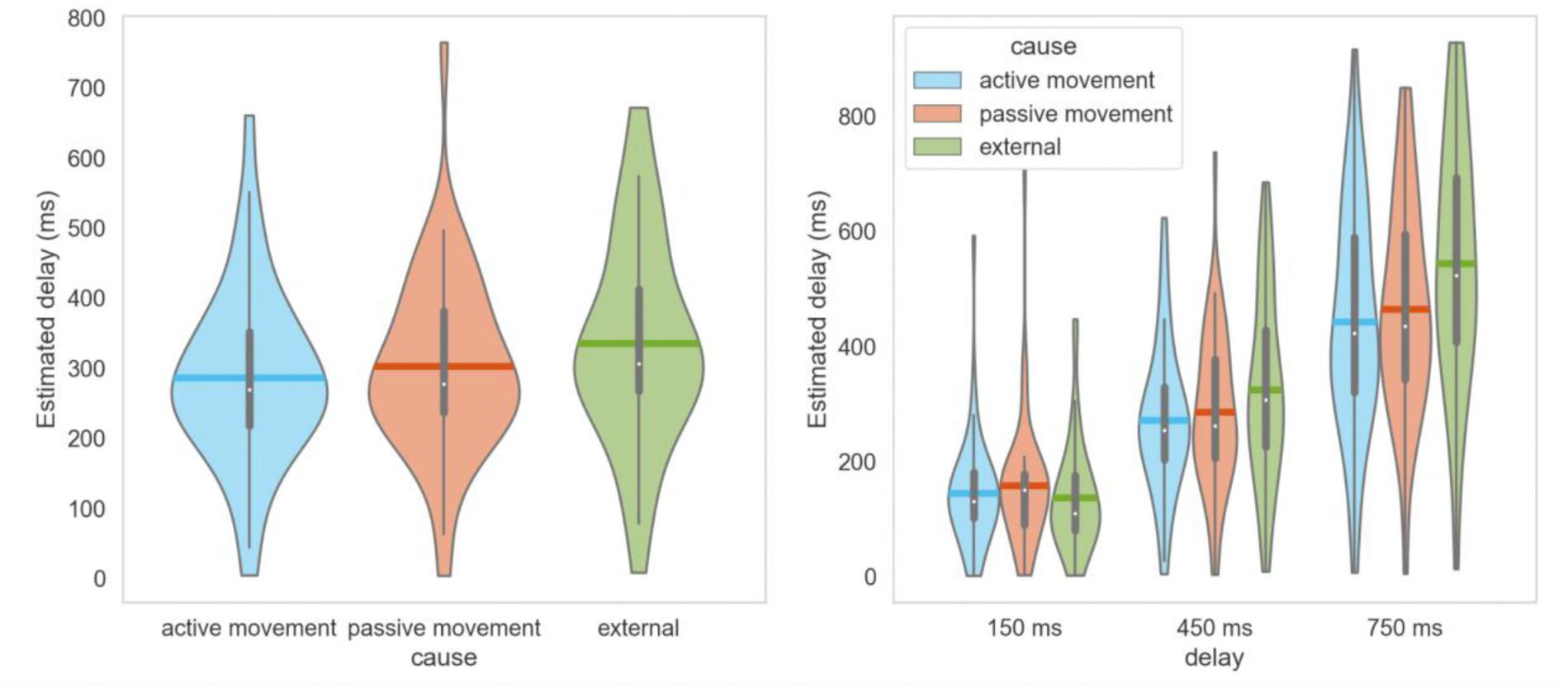
Results from Experiment 1 (auditory action outcome). **(A)** Average estimated outcome delay in the three conditions (active movement, passive movement and the external sensory event). **(B)** Same as a function of the actual interval. Vertical gray lines in each violin refer to the interquartile range and the horizontal lines refer to the means.

### Intermediate discussion

In Experiment 1, we failed to demonstrate the existence of intentional binding. While evidence in favor of non-intentional temporal binding was substantial, the evidence in favor of the intentional binding was inconclusive. Compared to previous studies, our experiment was not under-powered but appropriately controlled for potential confounding factors. In order to reproduce and generalize our findings to another sensory modality, we conducted Experiment 2, with a similarly large sample size and with a visual outcome instead of an auditory one.

## Experiment 2

### Method

#### Participants

For Experiment 2 (visual action outcome), we aimed for a similar sample size as Experiment 1 and recruited 45 participants. Two participants were excluded, one due to failure to produce temporal intervals varying monotonically with actual intervals and the other due to failure to comply with instructions to control the gamepad to report the interval estimations. The remaining 43 participants (24 females, age = 24.8 ± 4.4 years old) participated in all three conditions. Participants were naive as to the purpose of the study and reported normal or corrected-to-normal vision, normal audition and no neurological history. All participants provided informed consent before participation and received a payment for their participation. Procedures were approved by the ethics committee (CEEI/IRB Comité d’Evaluation Ethique de l’Inserm, n°21-772, IRB00003888, IORG0003254, FWA00005831) and adhered to the ethical standards of the Declaration of Helsinki except for registration in a database. Data was collected in 2021.

#### Apparatus and setup

The action consisted of a keypress on a gamepad, and participants viewed the visual scene including the fixation cross through a head-mounted display (HMD, Oculus rift CV1, resolution 2160 × 1200) in virtual reality (VR). The action outcome was a visual effect (the lightening of a light bulb, Figure 3A), which was presented at either of two task-irrelevant distances from participants (near: 45 cm, far: 4.5 m). The VR setup was chosen in order to test the impact of the spatial distance dimension on temporal binding. As this manipulation did not yield any difference in temporal binding, data from both distances were pooled together for analysis. The experiment was programmed using the unity platform (Unity 2018.4.22f1 Personal) and Microsoft Visual Studio 2019.

**Figure 3.**
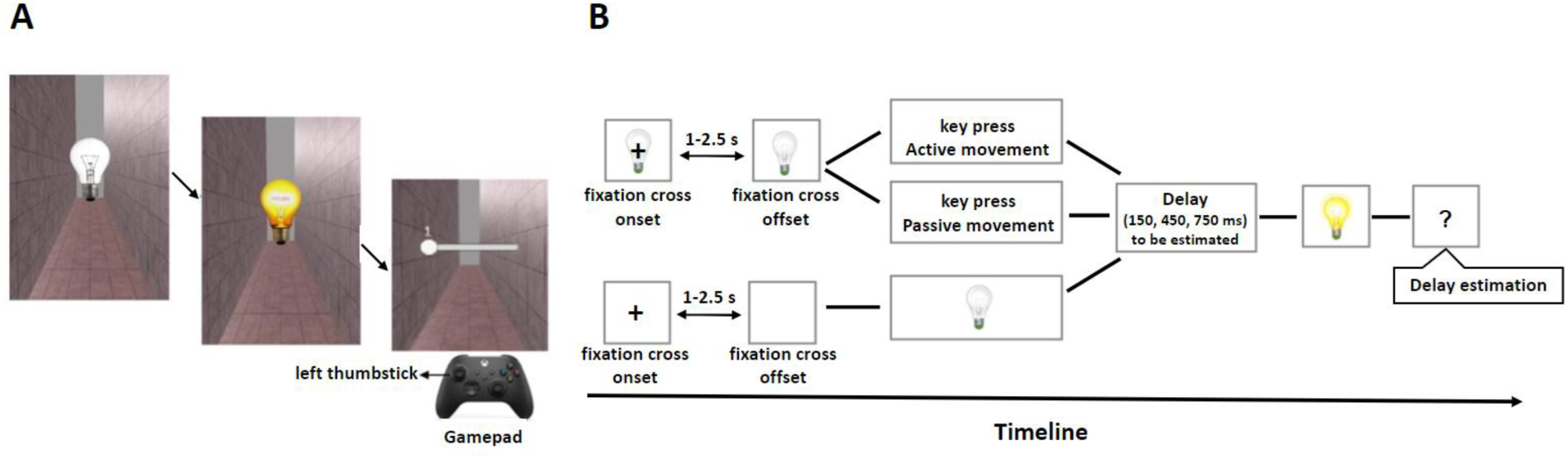
The experimental scene of a single trial (A) and experimental conditions (B) of Experiment 2. **(A)** During each trial, an off-bulb was turned on by different first events depending on the condition. The key-press and interval estimates were performed through a gamepad. Participants indicated their estimates by moving a visually depicted slider with the left thumbstick of the gamepad. **(B)** Schematic illustration of a trial in the three conditions: active movement, passive movement and external sensory event conditions. In the active and passive movement conditions, the light bulb (switched off) appeared at the beginning of each trial together with the fixation cross, while it appeared after the fixation cross in the external event condition. The light bulb remained visible prior to being illuminated. The lightened bulb lasted for 1 s before participants were asked to estimate the intervals. Note that the same number of events occurred during a trial for all the conditions.

#### Task and procedure

The procedure was similar to that of Experiment 1 except for two aspects: 1) the external sensory events were visual instead of auditory; 2) the key-press and interval estimates were performed through a gamepad. In the active and passive movement conditions, a light bulb was turned on once an active or a passive key-press (button “A” on the gamepad, Figure 3A) was performed. After the bulb had been lightened for 1 s, participants were then asked to estimate the interval between the onset of the key-press and the onset of the bulb lighting. In the external sensory event condition, participants merely viewed an off bulb being turned on by the computer. Participants were asked to estimate the interval between the appearance of the off bulb and its lighting onset. As participants were wearing the headset during Experiment 2, they indicated their estimates for all conditions by moving a visually depicted slider with their left thumb operating the stick of the gamepad (Figure 3A). The slider was presented to participants at a distance of 87 cm with a length of 54 cm and a width of 10 cm, and ranged from a minimum value of 1 on the far left to a maximum value of 1000 on the far right. Values appeared in real-time above the slider while participants moved it to the desired value, and they pressed the “LB” button on the gamepad to submit their responses. The intervals between the two events were the same as in Experiment 1 (Figure 3B) and the same number of repeated trials for each condition and for each interval were performed in Experiment 1 and 2. As in Experiment 1, the “Fix-off/First-event” duration was not directly recorded, but the “Trial-on/First-event” duration was recorded and shown to be similar for both the active and passive movement conditions (on average 2.67 s for the active and 2.77 s for the passive movement conditions, Mann-Whitney U test: p = 0.19). The distribution of the “Trial-on/First-event” duration in the external event condition did not mirror the trial-to-trial variability observed with the active and passive movement conditions (on average 2.23 s), but, again, allowed maintaining the predictability of the first event. Similar to Experiment 1, the timings for the external event conditions were different in a subgroup of 10 participants. For this reason, all the analyses were run again without this subgroup, and all statistical results that are different from the main analyses are reported as footnotes in the manuscript.

#### Data Analysis

The same analysis as for Experiment 1 was conducted.

## Results

There was substantial evidence in favor of the inexistence of a temporal binding when comparing the active movement condition (Mean ± SE = 245.15 ± 18.61) to the passive movement condition (Mean ± SE = 238.07 ± 18.02, BF_HN(0,122.5)_ = 0.15, RR_B<1/3_ = [57, >3000]). As in experiment 1, there was substantial evidence for the existence of a non-intentional binding when comparing the active movement condition to the external sensory event condition (Mean ± SE = 273.62 ± 18.11, BF_HN(0,122.5)_ = 3.3, RR_B>3_ = [7, 134]). Finally, there was substantial evidence in favor of a temporal binding when comparing the passive movement condition to the external sensory event condition (BF_U(13.5,122.5)_ = 62.8, RR_B>3_ = [0, 2274]).

When considering the delays separately, the evidence in favor of the inexistence of an intentional binding was inconclusive for the long interval (BF_HN(0,122.5)_ = 0.57, RR_1/3<B<3_ = [0, 214]) and substantial for the two other intervals (middle: BF_HN(0,122.5)_ = 0.25, RR_B<1/3_ = [93, >3000]; short: BF_HN(0,122.5)_ = 0.18, RR_B<1/3_ = [67, 3000])^5^. There was substantial evidence in favor of the existence of non-intentional temporal bindings for the longest interval (active vs. external sensory conditions: BF_HN(0,122.5)_ = 17.9, RR_B>3_ = [8, 810]); passive vs. external sensory condition: BF_U(13.5,122.5)_ = 18373, RR_B>3_ = [0, >3000]) and the middle interval (active vs. external sensory conditions: BF_HN(0,122.5)_ = 5.97, RR_B>3_ = [7, 252]); passive vs. external sensory condition: BF_U(13.5,122.5)_ = 657, RR_B>3_ = [0, >3000]). There was substantial evidence in favor of the inexistence of such bindings for the short interval (active vs. external sensory conditions: BF_HN(0,122.5)_ = 0.09, RR_B<1/3_ = [32, >3000]); passive vs. external sensory condition: BF_U(13.5,122.5)_ = 0.16, RR_B<1/3_ = [52, >3000]).

**Figure 4.**
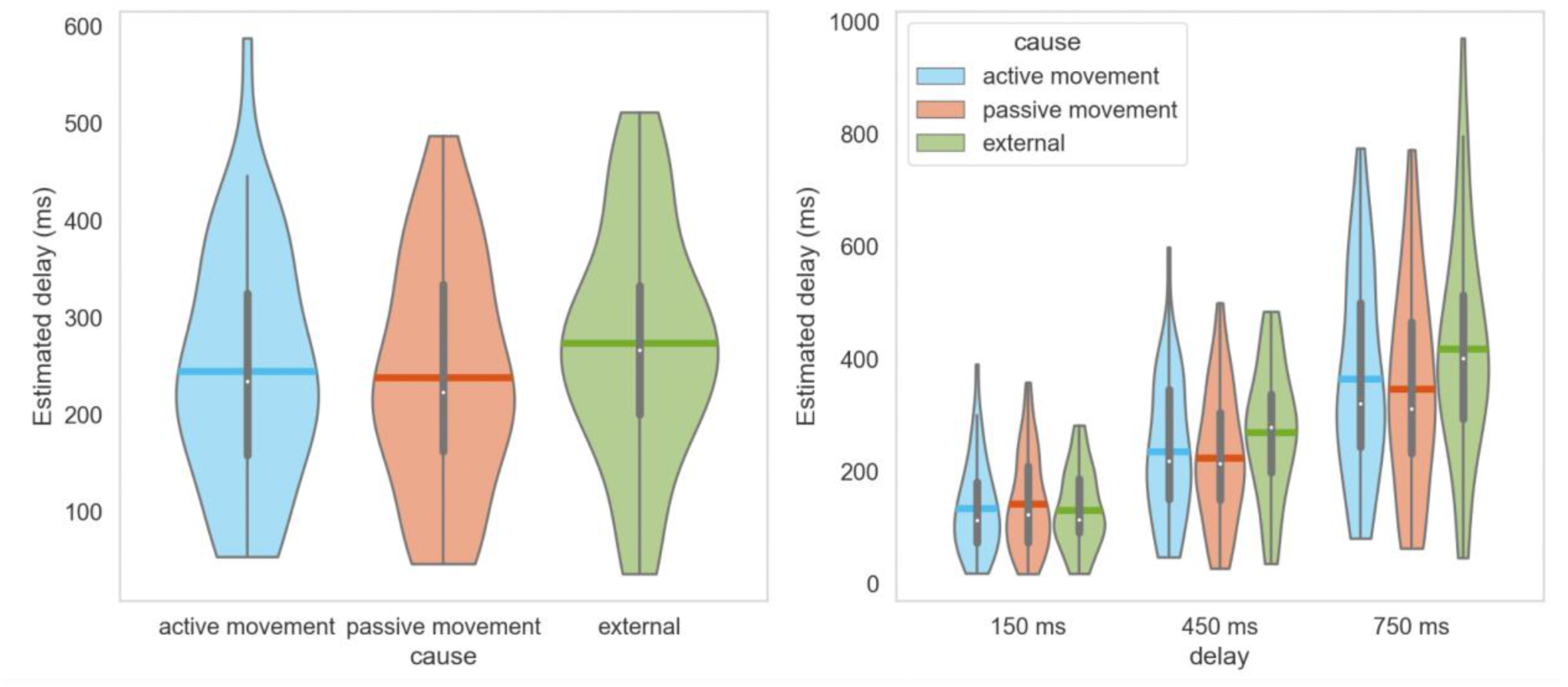
Results from Experiment 2 (visual action outcome). **(A)** Average estimated outcome delay in the three conditions (active movement, passive movement and the external sensory event). **(B)** Same as a function of the actual interval. Vertical gray lines in each violin refer to the interquartile range and the horizontal lines refer to the means.

### Intermediate discussion

Experiment 2 led to results similar to those observed in experiment 1, with even stronger evidence in favor of the inexistence of intentional bindings but supporting the existence of non-intentional bindings. The scope of these results will be discussed below.

### General Discussion

The present study examined whether the nature of temporal binding effect is genuinely intentional, while controlling for both the presence of somatosensory information and predictability of the first event, by comparing interval estimates between the active movement, passive movement and external sensory event conditions across two sensory modalities of the action outcome. In Experiment 2, we found that, when the outcome was visual, action intention was neither necessary for observing a binding (conclusive evidence for the existence of a non-intentional binding when comparing passive movement to the external sensory event conditions) nor sufficient (conclusive evidence for the inexistence of the intentional binding when comparing active to passive movement conditions). When the outcome was auditory, this evidence gathered from Experiment 1 was inconclusive (with Robustness Regions indicating that even smaller predictions of the size of the intentional binding would lead to the same conclusion). Thus, even with a larger number of participants than in previous studies with significant intentional binding but lacking appropriate controls, we could not demonstrate the existence of intentional binding in this experiment either. If an intentional binding existed with an auditory outcome, it would be in any case much weaker than previously assumed. However, we found a temporal (but not intentional) binding when comparing the passive movement condition to the external sensory event condition in both modalities that was modulated by the actual delay between the two events, with conclusive evidence for such binding mostly with the longest delay.

Here we carefully controlled for the available information between the active movement condition and the baselines. The difference of the available information for active and passive movements was that there was a voluntary motor command induced by intention in the active movement condition, but not in the passive movement condition. The absence of a substantial difference in temporal estimation between these two conditions suggests that action intention alone is not sufficient to induce intentional binding. This aligns with Buehner and Humphreys (2009) finding that intentional action that is not perceived as causally related to the second event is not sufficient for temporal binding to occur. Instead of the intentionality of an action, the ‘intentional’ binding observed in previous studies could be due to the uneven level of temporal predictability and/or causality for active and passive movements. Our results are also in keeping with a recent study showing that the binding difference between voluntary and involuntary movements, as measured with the Libet clock procedure, vanishes when the temporal predictability of to-be judged movements is controlled for (Kirsch et al., 2019). Our results go further by showing that intentional binding also vanishes with the interval estimation method, in both auditory and visual outcome modalities.

Interestingly, we further observed that the perceived temporal interval was shorter for both the active and the passive movement conditions, when compared to the external sensory event condition, in both experiments. This result indicates that the intentionality of an action is also not necessary for the temporal binding. This finding aligns with Suzuki et al. (2019); however, intention, predictability and proprioception differed between their compared conditions, whereas we varied only one piece of information between two compared conditions. This approach allowed us to disentangle the confounders of action intention, somatosensory information and predictability. Our results also align with Buehner (2015), and here we additionally control for the predictability of the first event. Compared to active and passive movements, the external sensory event condition lacks intentional efferent and somatosensory information about the action. It has been proposed that binding phenomena result from cue integration (Moore & Haggard, 2008; Moore & Fletcher, 2012; Moore et al., 2009; Wolpe et al., 2013), but this framework mainly relied so far on the role of efferent motor signals. The present findings stress and support the role of somatosensory information, in which efferent motor signals do not necessarily need to be present (Kirsch & Kunde, 2023; Kirsch et al., 2019). Of interest, somatosensory events conceivably have a fuzzier perceived time point than auditory or visual events, and the binding has been shown to be modulated by the reliability of the events to be bonded together (Klaffehn et al., 2021). Thus, the contribution of somatosensory information to the binding might be related to the less reliable temporal estimation of the somatosensory information in the active and passive movement conditions than of the visual or auditory first event in the external sensory event condition. Future studies, where the reliability of the baseline estimation is manipulated, will need to be conducted to explore this possibility.

It is worth noting that, when comparing between the external sensory event condition and the active/passive movement condition, the temporal attraction did not occur for the shortest 150-ms interval in either experiment. This observation is reminiscent of the hypothesis that temporal bindings might result from the causal relationship between two events, rather than from intentionality per se, as suggested by previous studies (Buehner, 2015; Buehner, 2012; Buehner & Humphreys, 2009; Desantis & Buehner, 2019; Desantis et al., 2011). Compared to longer delays in the present study, the first event and the effect are more likely to be perceived as causally linked in the shortest delay, as they follow each other closely in time. This could create a “causal” binding, i.e., a perception of the events closer than they really are, in all three conditions, strong enough to mask any additional binding revealed by comparing the conditions between them.

In sum, we did not observe intentional binding when comparing active vs. passive movements with the interval estimation procedure. Instead, temporal binding emerged when comparing both active and passive movements to an externally triggered sensory event. Taken together, these findings indicate that the link between the presence of a temporal binding effect and the intentionality of an action is neither necessary nor sufficient. Instead, temporal binding is not “intentional” and caution should be exercised when using this phenomenon as an implicit measure of the sense of agency. Our findings argue that intentional binding between voluntary and involuntary actions found by previous studies could be due to confounding factors and that experimental procedures such as the choice of the baseline, are crucial. These findings further stress the role of somatosensory integration and temporal prediction on the temporal attraction instead of motor command or action intention.

## Author Contributions

G.K. conceived and designed the study; G.K. programmed the study with the support from C.D.; G.K. and C.A. collected the data; G.K. analyzed the data with the support from M.V. and A.F.; G.K. wrote the paper; G.K., A.F., and M.V. revised the paper.

## Funding

This study was supported by the post-doctoral fellowship from Fondation Fyssen to G.K., it’s also supported by the ANR grant DEC-SPACE (ANR-21-CE28-0001), the Impulsion grant from IDEXLYON, and the LABEX CORTEX (ANR-11-LABX-0042).

## Acknowledgments

G.K. would like to thank her friend Shichun Hu for software assistance, Manon Petit for the help with data collection, Célia Farge, Sonia Alouche, Florence Tamissa-Leger and Sandra Chinel for administrative support and Eric Koun, Roméo Salemme for engineering support.

## Footnotes

1 With the group of 31 participants without technical error, there was substantial evidence for the inexistence of the intentional binding (BF_HN(0,122.5)_ = 0.22, RR_B<1/3_ = [83, >3000])

2 With the group of 31 participants without technical error, there was substantial evidence for the existence of such temporal binding (BFU(13.5,122.5) = 4.5, RRB>3 = [0, 163])

3 With the group of 31 participants without technical error, there was substantial evidence for the inexistence of the intentional binding for two of the intervals (long: BFHN(0,122.5) = 0.13, RRB<1/3 = [48, >3000]; short: BFHN(0,122.5) = 0.25, RRB<1/3 = [93, >3000])

4 With the group of 31 participants without technical error, there was substantial evidence for the existence of such temporal binding for middle interval (BFU(13.5,122.5) = 3.4, RRB>3 = [3, 124])

5 With the group of 33 participants without technical error, the evidence for the inexistence of the intentional binding was inconclusive for the middle interval (BFHN(0,122.5) = 0.54, RR1/3<B<3 = [0, 201])

## Notes

https://osf.io/n7y8a/

### Competing Interest Statement

The authors have declared no competing interest.

### Summary of Updates

Author Contributions clarified; Acknowledgements and Funding added; Title page updated.

